# In situ observation of water transport through the urothelium with optical coherence tomography

**DOI:** 10.1101/2025.03.03.641330

**Authors:** Lan Dao, Hui Zhu, Hui Wang

## Abstract

**Significance:** This study provides the first direct evidence of water transport through the urothelium, traditionally considered impermeable. Using Optical Coherence Tomography (OCT), we observe that the urothelium absorbs and expels water under varing concentrations of NaCl, challenging long-held views about its impermeability. The discovery that osmotic stress can induce urothelial damage has significant implications for bladder disorders like interstitial cystitis and overactive bladder, where urothelial integrity is compromised. These findings open new avenues for understanding the mechanisms behind these conditions.

**Aim:** Traditionally considered impermeable, the urothelium has recently been implicated in water transport due to the presence of aquaporins (AQPs). Despite this, direct evidence of water movement through the urothelium remains elusive. This study aims to provide such evidence by examining urothelial responses to NaCl solutions using Optical Coherence Tomography (OCT).

**Approach:** Fresh porcine bladder samples were subjected to OCT imaging to observe urothelial responses under varying osmolarity conditions, using NaCl solutions ranging from 0.31 Osm/L to 2.07 Osm/L. Urothelial thickness was measured pre- and post-NaCl application. Additionally, histological and scanning electron microscopy (SEM) analyses were conducted to assess cellular integrity and damage.

**Results:** OCT imaging revealed a significant increase in urothelial thickness following deionized water application, indicative of water absorption. Conversely, exposure to higher osmolarity NaCl solutions resulted in urothelial shrinkage, suggesting water efflux. Histological analysis demonstrated intact cellular structures at lower osmolarities (0.31 Osm/L) but significant cellular disruption at higher concentrations (≥1.03 Osm/L). SEM analysis corroborated these findings, showing progressive damage to umbrella cells with increasing osmolarity.

**Conclusions:** This study provides direct evidence that the urothelium is a dynamic barrier capable of water transport, modulated by osmotic gradients. The observed osmotic-induced urothelial damage may have significant implications for the pathophysiology of conditions such as interstitial cystitis and overactive bladder, offering new insights into potential diagnostic and therapeutic strategies.

---

The bladder is a round bag-like organ for storing urine and then expelling it under volitional control. The bladder wall consists of five layers of tissue, urothelium, lamina propria, submucosa, muscle, and fat. The urothelium is a thin layer lining the surface of the bladder wall formed by three cell layers, basal cell layer, intermediate cell layer, and umbrella cell layer. On the top of the umbrella cell layer, there is a thin glycosaminoglycan or GAG layer. The umbrella cells are closely packed together by tight junctions and covered with uroplakin, which provides strong protection for the urothelium^1^. The diffusive permeability of water and ions measured with transepithelial electrical resistance (TEER) and isotope tracer diffusion method seems very low^2,3^. Therefore, conventionally, the urothelium has been considered a physical barrier between the urine and the ‘blood’ to protect the underlying bladder tissues, against urine, toxins, and infections like bacteria.

The concept of the impermeability of the urothelium has been challenged from the very beginning of its development. Almost a century ago, an *in vivo* animal study demonstrated water and sodium chloride (NaCl) transport through the urothelium by tracking the concentration variations of the added hemoglobin and chloride ions^4^. In a later *in vivo* human study, water absorption was reported by tracking tritiated water^5^. With the advancements in molecular biology, researchers have discovered the expression of aquaporins (AQP), a family of water channel proteins, on the urothelium of a variety of species including humans. AQP-3, -4, -7, and -9 have been identified on human surgical tissue and differentiated normal human urothelial cells^6^; AQP-1, -2, and -3 have been identified on rat bladder, and AQP-1, -3, - 9, -11 have also identified on the porcine bladder^7,8^. These discoveries provided the molecular mechanism for water regulation and transport through the urothelium. With ultrasonic imaging, the bladder volume of a group of participants was monitored while sleeping^9,10^. Significant volume reduction of the urinary volume indicated water transportation in the bladder. The observation was further substantiated by monitoring the change in the concentration of a blue dye, which correlated with the alternation of bladder volume.

In the last century, various technologies have been employed to seek evidence if the urothelium is permeable. The microstructures shown on the umbrella cells seem to indicate that the urothelium is a barrier for protecting the underlying tissue, while the AQPs distributed through the urothelium suggest that the urothelium may regulate water or other ions transport. However, none of the demonstrated technologies can provide direct evidence of water transport through the urothelium. Optical coherence tomography (OCT) is a noninvasive imaging technology, which can acquire tissue cross-sectional images at a resolution of less than 10µm. OCT has been developed for urological cancer diagnosis as it can clearly differentiate the urothelium, the lamina propria, and the detrusor muscle^11,12^. In this report, using OCT, for the first time, we demonstrate *in situ* observation of the urothelial responses to NaCl solution under different osmatic concentrations due to water transport.

## MATERIALS AND METHODS

### OCT imaging

An in-house built spectral-domain OCT system was used for acquiring OCT images. A supercontinuum light source (YSL Photonics) filtered at ∼800nm with ∼150nm full width at half maximum was used as the broadband light source. The axial resolution is ∼2µm and the lateral resolution is ∼7.9 µm with a power of ∼2.3mW on the sample. OCT images were acquired at 17 frames per second with 1000 A-scans per frame. The raw data of the OCT images were then post-processed in Matlab for further analysis.

### Tissue preparation

Fresh porcine bladders were collected from a local slaughterhouse immediately after the animal sacrifice. The collected porcine bladders were transported and stored in Krebs-solution at 4°C. All experiments in this study were completed in six hours^13,14^. The bladder was first gently washed with deionized (DI) water, and then cut into four 3 cm x 3 cm specimens. The four edges of a tissue sample were mounted by clippers on four translational stages. During imaging, the tissue sample was gently and uniformly stretched by the translational stages from all edges so that the urothelium layer could be clearly visualized under OCT.

### Urothelial thickness measurement

A precision microscope cover slide (170µm±5µm, Thorlabs) was used to calibrate the thickness measurement based on OCT images. An OCT image of the cover slide was acquired with the identifiable bottom and top surfaces of the slide. The thickness per pixel (TPP) is calculated as

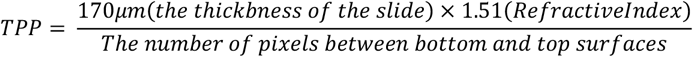

As the urothelial boundaries can be visualized in OCT images, the urothelial thickness can be calculated based on the number of pixels covered by the urothelial layer. On a single OCT image of a bladder tissue sample, the thickness at five different locations was measured and averaged as the thickness of the urothelium.

### Urothelial thickness variation

After a tissue sample was mounted, the tissue sample was first gently dried by blowing off the residual water on the surface with air for 10 seconds. Then, deionized water (DI) water was then added. OCT recorded the urothelial response while adding the DI water. For observing the urothelial response at different osmolarity concentrations, the tissue surface was first covered with a thin layer of DI water, and then 100μl 2.07Osml/L (6%) NaCl was added to the imaged site to replace the DI water.

### Urothelial thickness quantification at different osmolarity

After a tissue sample was mounted, 100μl of 0.31Osm/L (0.9%), 0.52Osm/L (1.5%), 0.69Osm/L (2%), 0.86Osm/L (2.5%), 1.03Osm/L (3%), 1.38Osm/L (4%), and 2.07Osm/L (6%) NaCl solution was gently and sequentially added to the surface of a tissue sample. The tissue surface was washed by DI water and the surface residual water was dried by a Kimwipe before adding the solution with different concentrations. The OCT recorded tissue responses through the solution-adding process. Seven tissue samples were quantified at each osmolarity.

### Urothelial damage at different osmolarity

A bladder was cut into six tissue samples and immersed into six NaCl solutions at 0Osm/L (water), 0.31Osm/L (0.9%), 0.69Osm/L(2%), 1.03Osm/L(3%), 1.38Osm/L(4%), and 2.07Osm/L(6%) for 10 minutes. Then, each tissue samples were cut into three pieces. Two pieces were fixed into 10% formalin for later standard hematoxylin and eosin (H&E) stain and another piece was fixed for scanning electron microscopical (SEM) image. For H&E, two histological slices were obtained from each tissue sample. For each concentration, 40 histological slides were obtained from 10 bladders.

## RESULTS

In the OCT images, the urothelium appears as a layer with relatively lower contrast atop the lamina propria, which exhibits stronger contrast. Since OCT contrast originates from backscattered light, it is directly related to the tissue’s scattering properties. Figure 1(a) and (b) illustrate the changes in urothelial thickness before and after the application of DI water to a dried urothelial surface. The thickness of the urothelium increased by approximately 52% following the addition of DI water, indicating significant water absorption by the urothelial cells. Figures 1(c) and (d) show the urothelial thickness before and after applying a 2.07 osm/L NaCl solution to a tissue sample. A notable shrinkage of the urothelium is observed between Figures 1(c) and (d), suggesting that intracellular water was expelled from the urothelial layer due to the osmolarity gradient. As a result of the water extraction, the cellular organelles within the urothelium were densely packed, enhancing light scattering observed under OCT. The arrow in Figure 1(d) points to a shadow beneath the urothelial layer, indicating that the intense scattering from the contracted urothelium obstructs the light backscattered from the lamina propria. The processes of the urothelial response can be watched in Video1 and Video2.

**Figure. 1.**
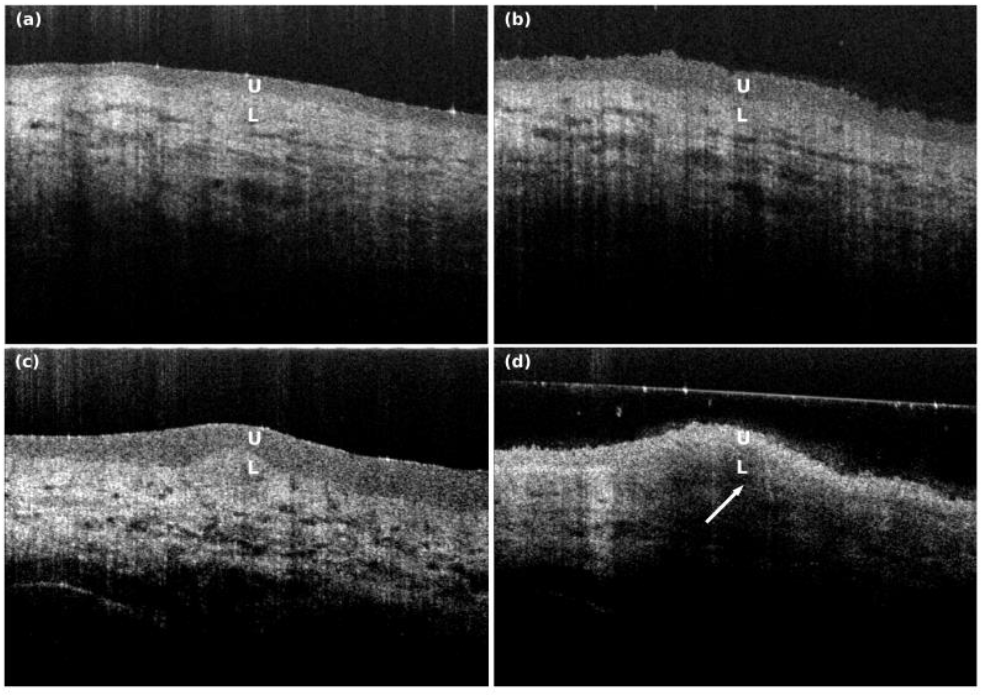
*In situ*. monitoring urothelial thickness change under OCT. (a) and (b) Urothelial thickness change before and after adding DI water; (c) and (d) Urothelial thickness change before and after adding 0.207 Osm/L NaCl solution. U: urothelium; L: Lamina propria (with video 1 and 2)

To quantify the urothelial responses treated with NaCl solution at different osmolarities, the urothelial thickness variation was measured using OCT images. As plotted in Figure 2(a), the urothelial thickness change is plotted against six discrete osmolarities. The experiment was repeated on seven tissue samples. Figure 2(b) further plots the percentage change in urothelial thickness in comparison to its initial thickness when treated with DI water. When the osmolarity is equivalent to normal saline (0.31Osm/L), urothelial thickness exhibits negligible changes from the baseline. However, as the osmolarity is increased to 0.52Osm/L, the thickness is decreased by approximately 20%. This contraction can reach around 30% at an osmolarity of 0.69Osm/L and escalate to 40% with a further increased osmolarity to 0.86Osm/L.

**Figure. 2.**
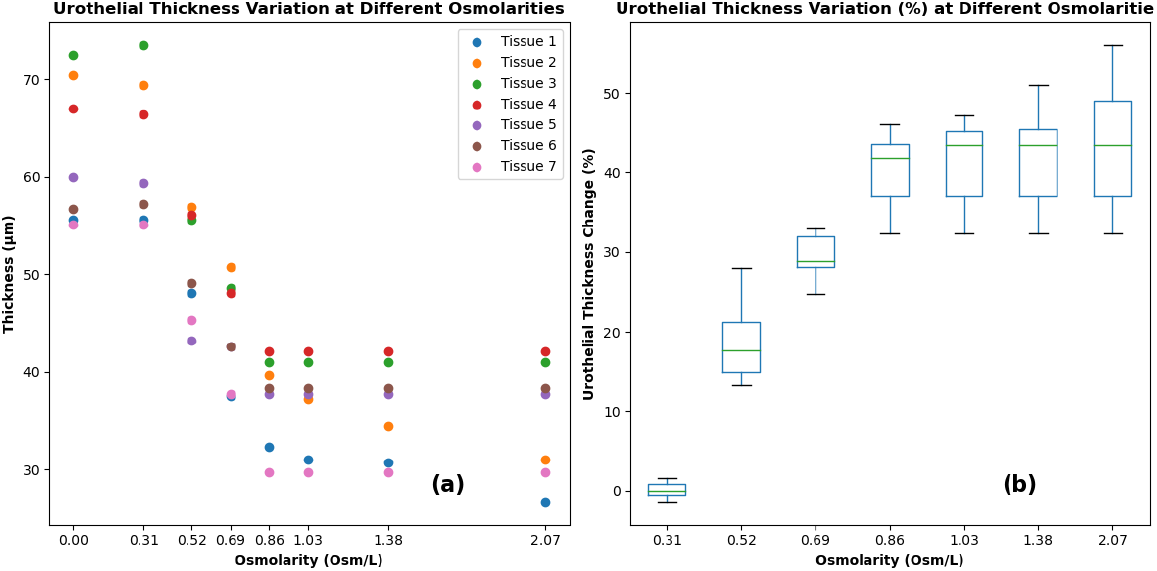
Urothelial thickness variation after applying NaCl solutions at different osmolarities. (a) The change of the urothelial thickness (µm) of seven tissue samples; (b) The percentage change of the urothelial thickness of seven tissue samples.

Beyond this concentration, further increasing NaCl osmolarity does not appear to induce additional urothelial shrinkage. The observed decrement in urothelial thickness is indicative of water efflux from the urothelium, presumably triggered by the osmotic gradient established between intracellular and extracellular environments.

Histological analysis was performed using H&E staining to assess cellular-level responses in tissue samples exposed to NaC solutions of varying osmolarities. The urothelium in samples treated with DI water and saline (0.31Osm/L) remained largely intact, displaying distinct layers including the umbrella cell layer, intermediate cell layer, and basal cell layer (Fig. 3a, Fig. 3b). Although the sample exposed to 0.69Osm/L NaCl exhibited an approximately 30% reduction in urothelial thickness (Fig. 2b), no overt damage was observable in the H&E image (Fig. 3c). More pronounced damage was noted in the sample treated with 1.03Osm/L NaCl, particularly in the umbrella cell layer, where discontinuities appeared on the surface of the urothelium (Fig. 3d). Increasing the NaCl concentration to 1.38Osm/L and 2.07Osm/L resulted in severe damage throughout the urothelial layers; at some locations, both the umbrella and intermediate cell layers were completely lost (Fig. 3e and Fig. 3f).

**Figure. 3.**
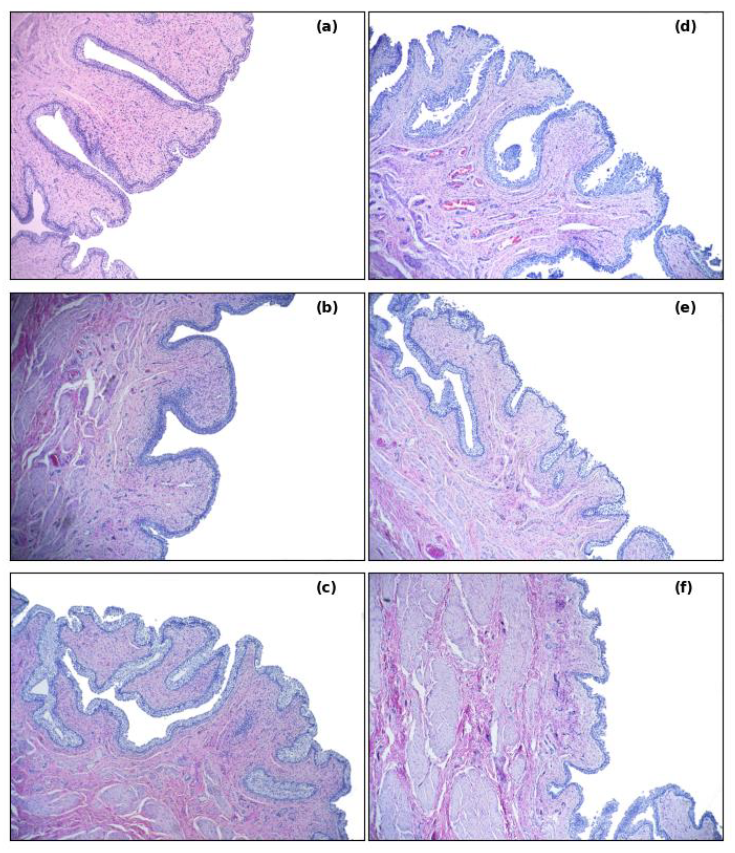
H&E (10×) of the tissue samples after applying NaCl with different osmolarities. (a) DI water. (b) 0.31Osm/L(Saline). (c) 0.69Osm/L. (d) 1.03Osm/L. (e) 1.38Osm/L. (f) 2.07osm/L.

Scanning electron microscopy (SEM) has been used to image the ultrastructure of the urothelial cells of the tissue samples treated with NaCl solutions. For the tissue samples treated with saline, intact dome-shaped umbrella cells covered with asymmetric unit membrane particles can be identified [Fig.4(a) and (b)]. This observation is aligned with the H&E image [Fig.3(b)], where the umbrella cell layer is complete. For the tissue sample treated with NaCl solution at 0.69Osm/L, the urothelium looks intact from the H&E image [Fig.3(c)]. However, on the SEM images [Fig.4(c) and (d)], it seems that the umbrella cells are still complete, but the surface of the umbrella cells has been subtly damaged. For the tissue sample treated with NaCl at 1.03Osm/L, the damage on the cell membrane of the umbrella cells is visible [Fig. 4(e) and (f)].

**Figure. 4.**
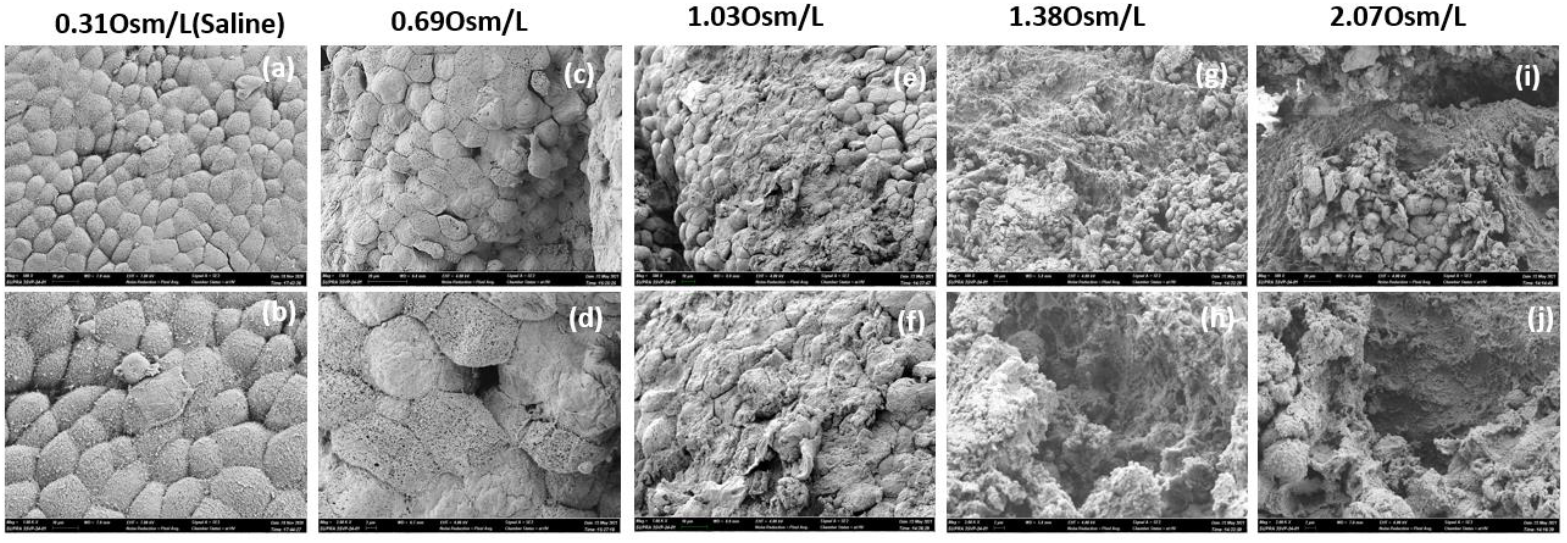
Scanning electron microscope images of the tissue samples treated with NaCl at different osmolarities. (a) and (b) Saline. (c) and (d) 0.069Osm/L. (e) and (f) 0.103Osm/L. (g) and (h)0.138Osm/L. (i) and (j) 0.207Osm/L.

Accordingly, such damage also appears in the histology image [Fig. 3(d)]. With the increased osmolarity, from SEM images shown in Fig. (h) to (j), severe damage to the urothelial layer can be observed. The loss of the entire umbrella cell layer exposes the below basal cell layers or even the lamina propria. A similar level of damage has also been observed in the corresponding histological images [Fig. 3(e) and (f)]. The SEM images confirm histology analysis observation and reveal some damage to the ultrastructure on the umbrella cells, which cannot be identified through histology.

## DISCUSSION

Historically, the urothelium has been examined through two primary perspectives: the impermeable barrier theory and the permeable barrier theory. The impermeable barrier theory posits that the urothelium functions as a robust physical barrier, preventing the passage of water, ions, and pathogens. This theory is supported by the structural integrity of umbrella cells, tight junctions, and the uroplakin layer, which together form a formidable defense against urinary components and pathogens. The permeable barrier theory, which has gained traction with advancements in molecular biology, suggests that the urothelium is selectively permeable and capable of regulating water and ion transport through specialized channels and proteins. Key among these are aquaporins (AQPs), a family of water channel proteins that facilitate the rapid transport of water across cell membranes. AQPs, such as AQP-3, -4, -7, and -9 in humans, have been identified in the urothelium, indicating a molecular basis for water permeability^6^. Additionally, *in vivo* observations of bladder volumetric changes during sleep provide further evidence that the bladder can regulate water^9,10^.

If water can be transported through the urothelium, the most direct evidence would be to observe the urothelial response to water transport. OCT, a non-tissue invasive imaging modality, enables us to visualize this process. Although the imaging depth of OCT is limited to 1-2 mm, its resolution can reach a few micrometers. In urology, OCT has been developed for the early diagnosis of bladder and urethral cancers^15,16^. When appropriately stretched, the bladder shows a layered structure where the urothelium, lamina propria, and muscularis layer can be visualized. Thus, OCT can be used to observe the urothelial response to water transport.

The *in situ* observation of the urothelial response to NaCl solution provides new insights into the permeability and structural integrity of the urothelium under varying osmotic conditions. Water transport is typically bidirectional. In Fig. 1(a) and Video1, the urothelial thickness increased by approximately 52% after the application of DI water over surface-dried tissue, indicating substantial water absorption by urothelial cells. The water absorption aligns with the observed reduction in bladder volume during sleep^9,10^. Conversely, exposure to high osmolarity NaCl solution (2.07osm/L) resulted in shrinkage of the urothelium. This osmotic challenge caused water efflux from the urothelial cells, leading to cellular dehydration and increased packing density of cellular organelles. The enhanced light scattering observed in OCT images, indicated by the shadow beneath the urothelial layer, underscores the structural changes occurring due to osmotic stress, as shown in Fig. 1(b) and Video2. Osmolarity appears to play a critical role in water transport^17^. In the second study, we conducted a quantitative analysis of urothelial thickness at varying osmolarities. While normal saline (0.31Osm/L) caused negligible changes, higher osmolarities led to significant contraction of the urothelium. The thickness reduction plateaued beyond 0.86Osm/L, suggesting that most of the free water had been expelled from the urothelial cells due to osmotic pressure.

Histological analysis allows us to observe changes in tissue microstructures due to significant water transport. The urothelial layer remains intact when treated with DI water and normal saline. However, increased osmolarity results in progressive damage, particularly in the umbrella cell layer. The damage is most pronounced at higher osmolarities, where both the umbrella and intermediate cell layers are severely affected. SEM images further confirm these observations, revealing ultrastructural damage not visible in histological images. The subtle damage to the umbrella cells at lower osmolarities and severe damage at higher osmolarities highlight the sensitivity of the urothelium to osmotic changes. The loss of the umbrella cell layer and exposure of underlying tissues at high osmolarities indicate significant disruption of the urothelial barrier. The damage to the urothelial cells could be related to osmotic lysis, when water exits the cells to balance the osmotic pressure in a short period time, the sudden cell shrinkage may lead to cell rupture^18^.

The findings from this study may have implications for conditions such as interstitial cystitis/painful bladder syndrome (IC/PBS) and overactive bladder (OAB). IC/PBS and OAB are complex disorders. IC/PBS is characterized by chronic pelvic pain, pressure, or discomfort associated with lower urinary tract symptoms, in the absence of infection or other identifiable causes, while OAB was defined as a syndrome as “urinary urgency, with or without urgency urinary incontinence, usually with increased daytime frequency and nocturia, if there is no proven infection or other obvious pathology.”^19,20^. The etiology of IC/PBS and OAB remains elusive, with various hypotheses proposed but none conclusively proven in clinical studies. For IC/PBS, a predominant theory suggests that urothelial damage induces bladder pain. This hypothesis assumes that compromised urothelial integrity increases permeability to concentrated urinary electrolytes, leading to inflammation and pain via activation of afferent sensory neurons in the bladder’s lamina propria.

For OAB, there is a theory hypothesis the urgency originated from the urothelium /suburothelium due to abnormal sensory function of the urothelium /suburothelium. However, it is not clear yet what causes urothelial damage.

In our study, with porcine bladder tissue, we observed damage to umbrella cells within minutes of exposure to NaCl solution. Previous studies have linked the permeability of urothelial cells to NaCl. It has been found that in differentiated human urothelial cells, AQP-3 is significantly upregulated by increased NaCl osmolality, but not by urea and glucose. In rats, AQP-2 expression on urothelial cells is significantly higher after injecting saline into the bladder compared to glucose. High salt intake has been shown to induce OAB-like symptoms in animal models^21–23^. Interestingly, clinical studies have demonstrated improved urinary symptoms in OAB patients following salt intake restriction^24^. These observations suggest that NaCl may play an important role in regulating water transport through urothelial cells, although the mechanisms have not been disclosed.

Under SEM, umbrella cell damage can be observed at an osmolality of 0.69Osm/L, a concentration that significantly exceeds typical sodium levels in human urine^25^. Our key finding is osmotic lysis resulting from substantial water transport. Beyond NaCl, AQP regulation in the urothelium is multifaceted. In rats, dehydration upregulates AQP-2 and AQP-3, bladder distension from mechanical stress increases AQP-2, and pathological conditions such as detrusor overactivity induced by partial bladder outlet obstruction upregulate AQP-1^8,26^. Additionally, factors such as vitamin D, intermittent fasting, and high-fat diets can modulate AQP-1 and AQP-3 expression^27^. Hormonal influences, particularly vasopressin, known to regulate AQP expression in renal tissues, may exert similar effects in the bladder^28^. The intricate regulation of AQPs suggests that significant water movement across the urothelium could occur if AQPs are upregulated due to various pathological conditions. Consequently, high urine osmolality might lead to urothelial cell damage or lysis. The diverse mechanisms of AQP upregulation could also account for the complex phenotypes observed in IC/PBS and OAB. However, this hypothesis requires rigorous validation through further research.

Creating controlled urothelial damage in animal models is also important for researchers to investigate physiological processes such as inflammation, cell death, regeneration, and scar formation in the bladder. Previously, such damages were induced by injecting toxic chemical agents (e.g., protamine sulfate) ^29,30^. Here, we show an easy and safe method to create progressive urothelial damage using NaCl solution at different osmolarities.

The present study has several limitations that warrant consideration. Primarily, all experiments were conducted *ex vivo* using porcine bladder tissue. While this model provides valuable insights, it may not fully replicate the complex physiological conditions present *in vivo*. To address this, large animal *in vivo* studies should be conducted to account for potential differences in tissue responses between *ex vivo* and *in vivo* conditions. Furthermore, these findings need to be confirmed using human bladder tissue, such as differentiated human urothelial cells. These additional studies would strengthen the translational potential of our observations and provide a potential solution for diagnosing and treating IC/BPS or OAB.

## Code and Data Avaliability

The raw data study will be provided upon request to the corresponding author.

## Acknowledgments

The U.S. Army Medical Research Acquisition Activity, 820 Chandler Street, Fort Detrick MD 21702-5014 is the awarding and administering acquisition office. This work was supported by the U.S. Army Medical Research Acquisition Activity, through the DoD PRMRP Discovery Award under Award No. W81XWH1810104. Opinions, interpretations, conclusions and recommendations are those of the author and are not necessarily endorsed by Department of Defense.

The paper is based on our previous SPIE Proceedings paper^31^.

## Notes

### Competing Interest Statement

The authors have declared no competing interest.

